# The effect of pulse shape in theta-burst stimulation: monophasic vs biphasic TMS

**DOI:** 10.1101/2023.03.06.531158

**Authors:** Karen Wendt, Majid Memarian Sorkhabi, Charlotte J. Stagg, Melanie K. Fleming, Timothy Denison, Jacinta O’Shea

**Author notes:** **Corresponding author:** Karen Wendt, Postal address: Institute of Biomedical Engineering, Old Road Campus Research Building, Oxford, UK OX3 7DQ. Currently with Magstim Ltd (Majid Memarian Sorkhabi). **Prior presentation** Abstracts on the early stages of this work were presented as posters at the annual meeting of the Society for Neuroscience (Neuroscience 2022) and the 5th International Brain Stimulation Conference. The manuscript has been posted on the pre-print server bioRxiv under the identifier https://doi.org/10.1101/2023.03.06.531158.

## Abstract

**Background:** Intermittent theta-burst stimulation (i)(TBS) is a transcranial magnetic stimulation (TMS) plasticity protocol. Conventionally, TBS is applied using biphasic pulses due to hardware limitations. However, monophasic pulses are hypothesised to recruit cortical neurons more selectively than biphasic pulses, thus yielding stronger plasticity effects. Monophasic and biphasic TBS can be generated using a custom-made pulse-width modulation-based TMS device (pTMS).

**Objective:** Using pTMS, we tested the hypothesis that monophasic iTBS would induce greater plasticity effects than biphasic, measured as induced changes in motor corticospinal excitability.

**Methods:** In a repeated-measures design, thirty healthy volunteers participated in three separate sessions, where monophasic and biphasic iTBS was applied to the primary motor cortex (M1 condition) or the vertex (control condition). Plasticity was quantified as changes in motor corticospinal excitability after versus before iTBS, by comparing peak-to-peak amplitudes of motor evoked potentials (MEP) measured at baseline and over 60 minutes after iTBS.

**Results:** Both monophasic and biphasic M1 iTBS led to significant increases in MEP amplitude. As predicted, monophasic iTBS induced a significantly larger effect than biphasic iTBS (linear mixed effect model analysis: (χ^2^(1) = 7.48, p = 0.006), which persisted even after subtracting each individual’s control (vertex) condition data from the M1 conditions (χ^2^(1) = 5.48, p = 0.019).

**Conclusions:** In this study, monophasic iTBS induced a stronger motor corticospinal excitability increase than biphasic within participants. This greater physiological effect suggests that monophasic iTBS may also have potential for greater functional impact, of interest for future fundamental and clinical applications of TBS.

## Introduction

Transcranial magnetic stimulation (TMS) is a non-invasive tool for neuroscientific research and is increasingly used for diagnosis and therapy in clinical practice [1]. It uses the fundamental principles of magnetic induction to modulate the nervous system: a brief electric current is applied to a stimulating coil, which creates a rapidly changing magnetic field that induces a voltage in the brain tissue underneath the coil. When applied repeatedly, TMS can induce plasticity – causing a change in cortical excitability of the targeted brain area that outlasts the stimulation period [2].

Different stimulation waveforms have been shown to recruit different neural populations, have different excitation thresholds, and have different effects on corticospinal excitability [3-8]. However, the range of stimulation pulses and patterns that can be generated by conventional TMS devices is limited by the device hardware and is usually confined to either monophasic or biphasic damped cosine pulses, where the exact shape and length of the pulse is determined by the resonance between the device components [9]. For monophasic pulses, the current flow is commonly stopped half-way through the cycle of the cosine pulse and the energy is dissipated through a resistor. This restricts not only the choice of TMS pulse waveforms and widths but also the achievable repetition rates [10]. For example, one class of repetitive TMS protocol, widely used for plasticity induction in fundamental research and clinical applications, that is constrained by these hardware limitations, is theta burst stimulation (TBS). During TBS bursts of 3 pulses are applied at 50 Hz and repeated every 200 ms [11]. In intermittent (i)TBS, a largely excitatory protocol, these triplets are applied for 2 s followed by an 8 s break and then repeated again, for 600 pulses in total [11]. To sustain these repetition rates, large amounts of energy need to be recovered after each stimulation pulse, and so TBS can usually only be delivered via a conventional TMS device using biphasic stimulation pulses.

Monophasic pulses are thought to more selectively recruit cortical neurons and have been shown to more strongly modulate cortical excitability than biphasic pulses when used in other repetitive TMS protocols [1, 6, 8, 12, 13]. For example, in quadripulse stimulation (QPS), bursts of four pulses are applied at inter-stimulus intervals of 1.5-1250 ms, repeated every 5s over 30 min [14]. A study comparing the after-effects of monophasic and biphasic QPS found that monophasic QPS induced stronger and longer lasting after-effects compared with biphasic QPS [13]. Such findings lead to the hypothesis that applying TBS with monophasic pulses may be more effective than existing biphasic TBS.

Recent technological developments of TMS devices using switching circuits, rather than the conventional resonance circuits, have allowed more control over TMS parameters and better energy recovery from the stimulation pulses [9, 15-17]. The programmable (p)TMS, a TMS device developed within our research group, which uses pulse-width modulation (PWM) to control cascaded inverters, enables more control over the pulse shapes by approximating a reference pulse of arbitrary shape using discrete voltage levels [9]. Previous evidence from computational modelling and an in-human physiology study indicated that the approximations of conventional pulse shapes generated using the pTMS have similar effects on the motor corticospinal excitability of healthy volunteers as the pulses generated by a conventional TMS device [18, 19]. Additionally, the pTMS device recovers energy effectively after each pulse, making the generation of monophasic TBS possible.

In this study, we use the pTMS device to generate monophasic and biphasic TBS and compared the effects on motor corticospinal excitability of healthy volunteers. We predicted that monophasic TBS would produce a larger plasticity effect (higher MEP amplitudes) than biphasic TBS. To control for intra- and inter-individual variability, we also applied the same stimulation to the vertex (control) and subtracted each individual’s MEP time-course in that control condition from each of the two active (motor cortex) TBS conditions, to test if the predicted difference would still be upheld.

## Materials and Methods

### Ethical approval

This study and the use of the pTMS device in this study were approved by the local ethics committee at the University of Oxford (Central University Research Ethics Committee, R75180/RE008). All participants gave their written informed consent prior to participating and were compensated for their time with £10/hr.

### Participants

30 healthy volunteers (16 females, aged 19-33 years, mean age 24.5 years) participated in one familiarisation session followed by three data collection sessions for this single-blind, within-participants crossover study. All participants were right-handed as assessed by the Edinburgh Handedness Inventory [20]. Participants were screened to rule out any current significant medical condition and any contraindication to TMS in line with international safety guidelines [21].

### Transcranial magnetic stimulation

The TBS intervention protocols were applied using the pTMS stimulator. The pulse waveforms generated by the pTMS stimulator were designed to closely approximate the conventional biphasic and monophasic pulses generated by a Magstim Rapid^2^ and a Magstim 200, respectively (see [18] for a detailed comparison). To measure the motor corticospinal excitability before and after TBS, a Magstim 200 stimulator (Magstim Co., UK) was used to generate monophasic single-pulse TMS to induce MEPs. A 70 mm figure-of-8 coil (Magstim Co., P/N 9925–00) was used to deliver all stimulation. Owing to coil overheating, for participants with resting motor thresholds (RMTs) above 43% of the maximum stimulator output (MSO) of the Magstim 200 (N = 3), one coil was used for MEP measurement and a second coil for TBS. For all other participants, the same coil was used throughout.

Prior to the three test sessions, there was an initial familiarisation session for participants naïve to TMS, where TMS was introduced to the participant and the hotspot and thresholds for the different parameters and devices were found. The pTMS stimulator’s maximum pulse amplitude is 1600 V, compared to the maximum amplitude of the Magstim Rapid^2^ and Magstim 200, which are approximately 1650 and 2800 V, respectively [19]. Therefore, to ensure the pTMS stimulator could generate iTBS at 70% of the RMT for both monophasic and biphasic pulses, individuals with RMTs above 47% MSO of the Magstim 200 were excluded from any further participation in the study (N = 3).

During the familiarisation and test sessions, participants were seated in a chair with their arms resting on a pillow on top of a table in front of them. The ‘motor hotspot’ of the left primary motor cortex was defined as the scalp location over which the lowest TMS pulse intensity elicited MEPs in the relaxed first dorsal interosseous (FDI) muscle of the right hand. For all TMS pulses, the coil was held by the operator and oriented at 45º to the midline with the handle pointing posteriorly, which results in a posterior-anterior current flow in the brain for the monophasic pulse. The direction of the biphasic pulse was reversed via the software, such that the direction of the dominant second phase of the pulse matched the current flow of the monophasic pulse (Fig. 1) [3, 5]. A Brainsight neuronavigation system (Rogue Research Inc., Montreal, Canada) was used to record the motor hotspot and for continuous tracking to maintain the position and orientation of the coil. Surface electromyography (EMG) of the right FDI was recorded in a belly-tendon montage (see supplementary file for details).

**Fig. 1:**
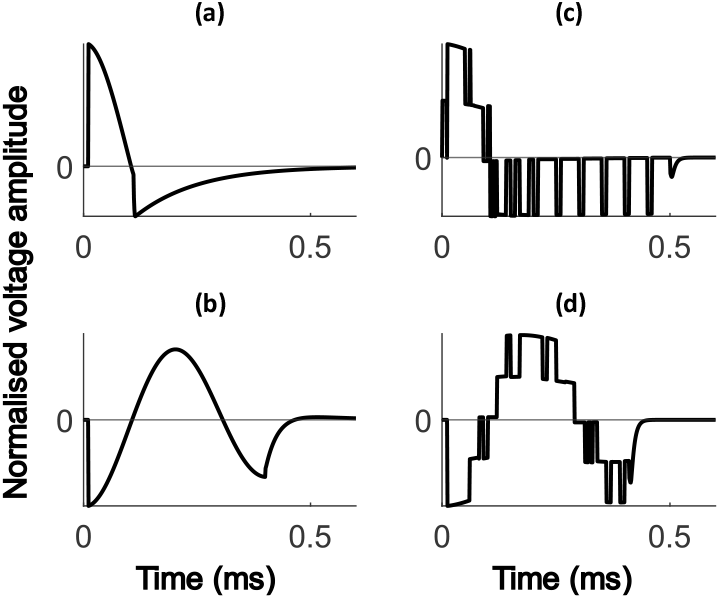
Simulations of the Magstim and pTMS pulse waveforms. Normalised simulated voltage waveforms of **(a)** the Magstim 200 and **(b)** the Magstim Rapid^2^ stimulators which were used as the references pulses to generate **(c)** the monophasic pulses and **(d)** the biphasic pulses with the pTMS stimulator. The direction of the biphasic pulses in (b) and (d) was adjusted such that the dominant second phase of the pulse matched the direction of the monophasic pulse.

The RMT, defined as the minimum intensity required to evoke an MEP of ≥ 50 μV peak-to-peak amplitude in 5 out of 10 consecutive trials, was determined by applying 10 pulses at each intensity and inspecting the EMG traces visually in real time for each device and pulse shape. To find the RMT, the pulses were triggered automatically via scripts in Signal version 7.01 (Magstim device) and Control desk (pTMS device) software at inter-pulse intervals of 5 seconds (± 15%).

Baseline excitability before and after TBS was quantified by blocks of 30 single TMS pulses at 120% of the RMT at inter-pulse intervals of 5 seconds (± 15%). The TBS protocol consisted of 600 either monophasic or biphasic pulses applied at 70% of the RMT [22].

### Procedure

The familiarisation and data collection sessions were at least one week apart, with each participant’s total duration of participation not exceeding 10 hours. During each data collection session the timeline was as follows (Fig. 2). After confirmation of the hotspot and the motor threshold, two baseline blocks of MEPs were recorded 5 minutes apart (30 pulses per block). iTBS was applied 10 minutes after the start of the first baseline block and follow-up blocks were recorded every 5-10 minutes after the start of the TBS protocol over the following hour (at 5, 10, 15, 20, 30, 40, 50 and 60 min). For each participant, two sessions were M1 conditions, where TBS was applied to the motor hotspot, and one session was a control condition, where TBS was applied to the vertex (similar to [23]). In the control condition, participants were randomized to receive either monophasic TBS (N = 14) or biphasic TBS (N = 16). The coil was lifted from the participants’ heads between each stimulation block and participants were instructed to keep their hands as relaxed as possible throughout.

**Fig. 2:**
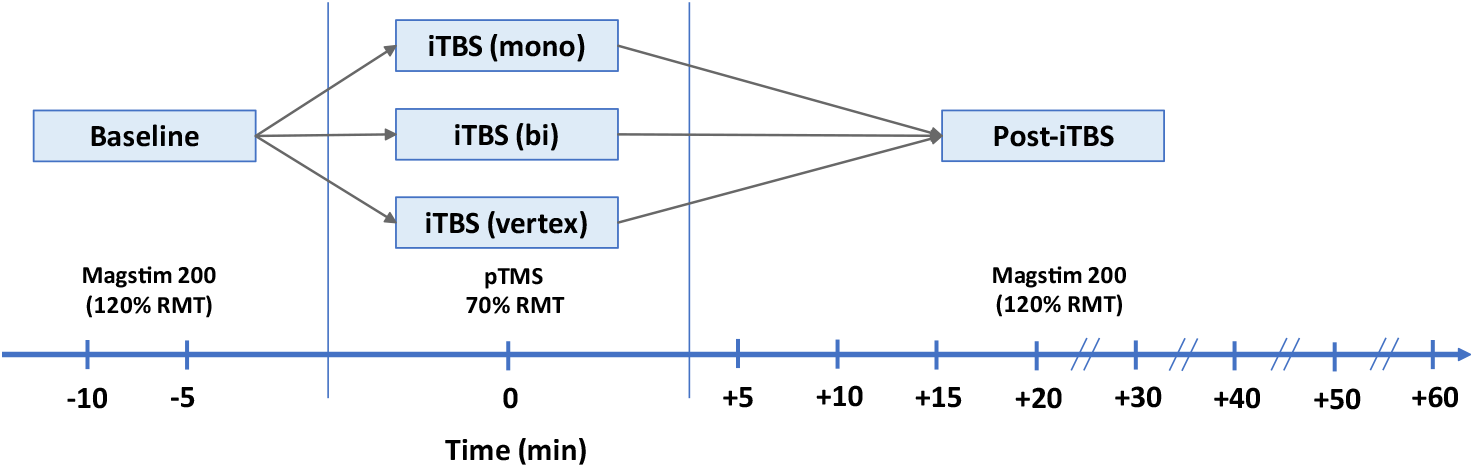
Overview of study design representing the flow of data collection at each visit. Each participant received each TBS condition on separate days (at least one week apart) in counter balanced order. Baseline MEPs in response to single-pulse TMS (30 pulses applied at 120% of the resting motor threshold) were collected in 2 separate blocks 10 and 5 min before the start of the TBS administration. iTBS was applied to the primary motor cortex (M1 condition) or the vertex (control condition) using monophasic or biphasic pulses for 190s. After TBS, MEPs were collected every 5 min for the first 20 min and then at 10-min intervals up to 60 min post-TBS.

The three test sessions were performed at the same time of day for each participant and the session order was randomised and counterbalanced. The participants were blinded to the stimulation condition.

### Data processing

The data were processed in Python using custom scripts. Since muscle activation can influence MEP amplitude, trials were excluded if the root mean square (RMS) of the EMG trace in the 90ms before the TMS stimulus (excluding the 5ms preceding the pulse) exceeded 0.02 mV. The EMG recordings included burst noise, a type of electrical noise characterised by sudden transitions between discrete voltage levels. To distinguish between this noise and any muscle activation, the pre-stimulus RMS was compared to the RMS during the silent period after the MEP (60 – 90 ms after the TMS pulse), where no muscle activity is expected but electrical noise may be present. Any trials where the pre-stimulus RMS was 0.005 mV larger than the RMS of the silent period or where the MEP was smaller than the peak-to-peak amplitude of the electrical noise (0.039 mV) were also excluded. The peak-to-peak MEP amplitude was calculated in the 15 – 60 ms time window after the TMS pulse was applied. Since the MEP amplitude data in over 55% of the stimulation blocks were not normally distributed (Shapiro-Wilk < 0.05), for each time point block the median (rather than the mean) MEP amplitude was calculated without applying any outlier removal. The medians from the two baseline blocks were averaged for each participant to give one baseline score per session. For each participant within each condition, the post-TBS change in MEP amplitude was calculated by subtracting the baseline for that condition from each post-TBS time point block, similar to McCalley et al [24]. Analyses were performed on these MEP change data. In addition, grand-average MEP change data were calculated for each condition by averaging over all post-TBS blocks (5-60 mins) and then contrasting across conditions. For time-course analyses, the vertex data of each individual were subtracted from the M1 TBS conditions for each of the post-TBS timepoint blocks, to control for inter- and intra-individual variability in MEP amplitude.

### Data analysis

To test for significant differences in pre-iTBS baseline excitability, repeated measures analysis of variance (rmANOVA) with factors Time (baseline blocks 1 and 2) and Condition (monophasic, biphasic, control) was used to compare the peak-to-peak MEP amplitudes of the two baseline blocks within and across conditions. Resting motor thresholds across testing sessions were also compared using rmANOVA with the factor Condition (monophasic, biphasic, control).

To investigate whether each iTBS condition induced plasticity (i.e. a significant increase in MEP amplitude), the data were averaged over all post-TBS time points and subtracted from the baseline to yield a single change score per participant per condition. These grand-average MEP change data were then compared against zero using one-sample t-tests. To test the *a priori* one-way directional prediction that both of the M1 stimulation conditions (monophasic and biphasic TBS) would induce plasticity (i.e. *increased* MEP amplitude), a one-tailed test was used. For the control (vertex) condition a two-tailed test was used. To compare relative plasticity induction across conditions, pairwise one-tailed t-tests on these change scores were conducted according to the *a priori* prediction of larger plasticity induction for monophasic than biphasic TBS and for M1 stimulation than vertex stimulation. Bonferroni correction for multiple comparisons was applied. Effect sizes are reported using Cohen’s d for one-sample t-tests and Cohen’s dz for pairwise t-tests [25].

To make use of the full MEP time-course data and to directly compare the effect of pulse shape when stimulating the motor cortex, complementary analyses were also run on the M1 conditions (monophasic or biphasic motor cortex stimulation) using linear mixed effects (LME) models. One advantage of this approach over rmANOVA is that it enables the inter- and intra-participant variability in the baseline data to be modelled in the analysis, as opposed to accounting for the baseline variability by calculating MEP percentage change scores, an approach which often fails to correctly model physiological processes [26]. In contrast to the previous grand-average MEP change analysis, the MEP change for each post-TBS time point were used, without grand averaging over the time points. In the LME models, baseline MEP amplitude, time (5-60 min post-iTBS) and TBS condition (monophasic, biphasic) were modelled as fixed effects while participants were modelled as a random effect. This allowed model intercepts to differ for different participants. To test the hypothesis that monophasic M1 TBS would induce stronger plasticity than biphasic, likelihood ratio testing was used to contrast two models – one that included TBS condition as a factor in the model versus a model without the TBS condition. The χ^2^ statistics representing the difference in deviance between the two models are reported, together with the p values calculated by the *anova* function using the Satterthwaite’s method for denominator degrees-of-freedom and F-statistic [27]. The analysis was repeated with the vertex control-subtracted data to account for inter- and intra-individual variability in MEP amplitude across post-TBS time points. All linear mixed effects models were created and analysed using purpose-written R code using the LME4 and lmerTest packages [27, 28]. The significance level was set to 0.05 for all analyses.

## Results

### No differences in RMTs or MEP amplitude at baseline

Resting motor threshold intensities did not differ between conditions (F(2, 58) = 0.43; p = 0.65). Also, a two-way rmANOVA showed that within sessions there were no significant differences between the first and second baseline measurements (F(1, 29) = 1.18; p = 0.29), nor did these differ across the TBS sessions (F(2, 58) = 0.11; p = 0.89; Supplementary Fig. S1). Thus, these analyses confirmed that participants were tested at comparable levels of motor corticospinal excitability prior to TBS in all three conditions.

### Both active iTBS conditions led to increased motor corticospinal excitability

Fig. 3 shows the group mean average change in MEP amplitude over the follow-up period (5-60 min post-TBS) after the baseline mean for each participant has been subtracted. As predicted, biphasic iTBS induced a significant increase in MEP amplitude (t(29) = 3.67, p < 0.001; Δ<u>M</u>: 0.19mV, SEM: ± 0.05mV; d = 0.67), confirming a plasticity effect. Monophasic iTBS also induced significant plasticity (t(29) = 4.53, p < 0.001; Δ<u>M</u>: 0.30 mV, SEM: ± 0.07mV; d = 0.83). By contrast, in the control condition, iTBS over the vertex did not lead to a significant MEP increase (t(29) = 1.30, p = 0.204; Δ<u>M</u>: 0.06mV, SEM: ± 0.05mV; d = 0.24).

**Fig. 3:**
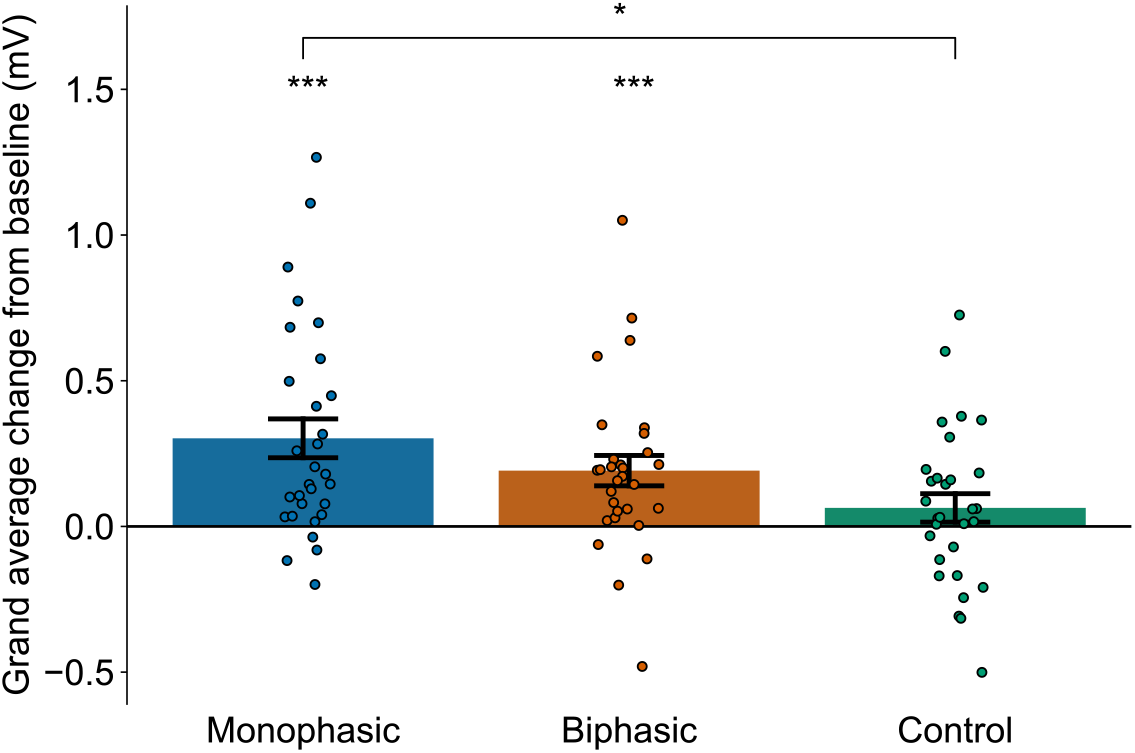
Group mean grand-average change in MEP amplitude compared to baseline across 60-min post-iTBS time period for the M1 (monophasic and biphasic) condition and the control (vertex) condition. Monophasic and biphasic iTBS led to significant increases in MEP amplitude. Only the monophasic plasticity effect was significantly larger than the control condition. Individual participants are indicated by dots, the bars indicate group means and the error bars represent +/-1 standard error of the mean. Asterisks indicate significance (***: p < 0.001; *: p < 0.05).

Bonferroni corrected pairwise one-sided t-tests showed that only the monophasic plasticity effect was significantly larger than the control condition (t(29) = 2.82; p = 0.013; dz = 0.52; monophasic vs biphasic: t(29) = 1.59; p = 0.183; dz = 0.29; biphasic vs control: t(29) = 2.07; p = 0.071; dz = 0.38). This analysis indicates that both monophasic and biphasic M1 iTBS induced plasticity, with only the monophasic iTBS leading to larger plasticity induction than the control condition when averaging over post-TBS time points.

### TMS pulse shape affects the iTBS plasticity effect

Fig. 4a shows the full time-course of changes in MEP amplitude across the 60-minute follow-up period for each stimulation condition. Raw plots (of non-baseline subtracted data) for each individual participant, timepoint and condition are shown in Supplementary Fig. S2. To directly compare the effect of pulse shape in the M1 conditions, the LME models with and without the fixed effect of TBS condition were compared using likelihood ratio testing, which showed that the pulse shape during iTBS (monophasic, biphasic) had a significant effect on the MEP change (χ^2^(1) = 7.48, p = 0.006). Fig. 4b shows the model fit of the LME model including the fixed effect of TBS condition for both M1 conditions. In line with the findings from the grand average analyses (Fig. 3), the summary output for the LME model, which includes the factor TBS condition, indicated that the MEP change was on average 0.11 mV higher in the monophasic relative to the biphasic TBS condition (B = 0.11, SE = 0.04, t = 2.75). Similarly, after subtracting the MEP values of the control condition from the M1 conditions, to control for the variability across the post-TBS time window (Fig. 4c), the likelihood ratio test still showed a significant effect of the pulse shape on the change in MEP amplitude (χ^2^(1) = 5.48, p = 0.019), with changes being on average 0.11 mV higher in the monophasic versus biphasic TBS condition (B = 0.11, SE = 0.05, t = 2.35). This analysis confirms that when considering the full MEP time-course data, monophasic iTBS induced a stronger motor corticospinal excitability increase than biphasic iTBS, even when subtracting the control condition to account for inter- and intra-individual variability in MEP amplitudes.

**Fig. 4:**
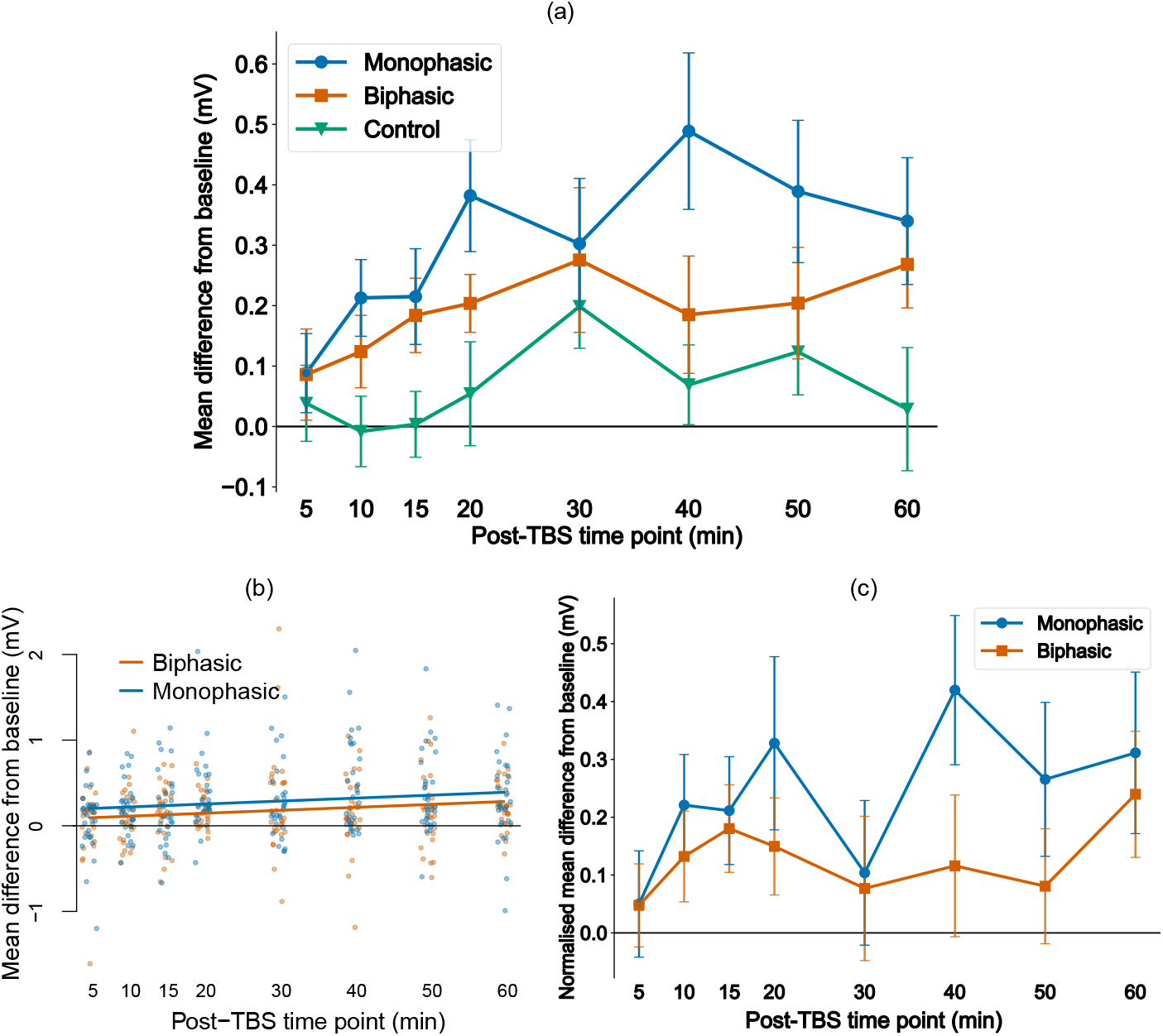
Group mean change in motor evoked potential amplitude (difference from baseline) over time for the different TBS conditions. (a) The mean change in MEP amplitude for the monophasic and biphasic M1 iTBS conditions and the control condition are shown across the post-TBS time points. (b) Visualisation of the fit of the linear mixed effect models to the data from the M1 (monophasic and biphasic) TBS conditions. Solid lines show the model prediction, single dots show partial residuals as generated using the ‘visreg’ function in R. (c) The group mean MEP change data over time for the monophasic and biphasic M1 TBS conditions, normalised by the control condition by subtracting the vertex change data for each participant across all post-TBS time points. Error bars indicate standard error of the mean.

## Discussion

In this study, both monophasic and biphasic iTBS applied to the primary motor cortex increased motor corticospinal excitability as measured using MEPs. The application of iTBS using the pulse-width modulation based TMS device resulted in excitability increases, as expected from studies applying iTBS using conventional resonance-based devices. On the group level, monophasic iTBS induced larger plasticity effects than conventional biphasic iTBS, confirming the importance of pulse shape in repetitive TMS protocols. TBS applied to the vertex in the control condition did not lead to significant MEP increases and the effect of pulse shape in the M1 conditions was sustained even when subtracting the control condition data from the M1 conditions.

### Choice of probing pulses

To measure the changes in motor corticospinal excitability, single pulses were applied at 120% of the RMT, similar to previous work such as [29, 30]. Other studies, including the early TBS studies [31], did not use % of RMT as a baseline/probe, but instead used the individualized intensity of TMS needed to reliably elicit MEPs of 1mV on each trial. However, this approach can suffer from floor or ceiling effects across individuals (e.g. in some participants, TMS may not induce peak-to-peak amplitudes much higher than 1mV) which can contribute to the high variability across participants [32]. Therefore, plasticity effects have been suggested to best be probed at a percentage of the resting motor threshold to take into account this difference in the input-output characteristics of the participants [33]. This reduces the risk of ceiling and flooring effects on plasticity but entails higher variability between participants at baseline. This was accounted for in the analysis of this study by including the baseline as a factor in the LME model.

Consistent with the literature, monophasic single pulses applied using a conventional Magstim 200 were used to measure MEPs pre/post iTBS. While this allowed the direct comparison of effects across conditions, monophasic and biphasic pulses may activate different neural populations during TBS. Using monophasic pulses therefore limits the ability to probe the potentially different neural populations activated by biphasic TBS [12, 34]. However, a study using both monophasic and biphasic pulses to assess plasticity effects found the results of using both pulse shapes highly comparable [35].

### Directionality of pulse currents

The direction of the current induced in the brain affects which neuron populations are activated and influences the size of the motor threshold [3, 5]. In this study, the monophasic pulses were applied to induce currents in the posterior-anterior direction both for single pulses and the TBS protocol, as this has been shown to have the lowest thresholds [5] and may therefore be the best current direction to increase motor corticospinal excitability. The biphasic pulses were applied to match this current direction in the second phase of the pulse, as this is thought to be the dominant activating phase of the biphasic pulse [5]. Future studies should evaluate and compare the effects of different current directions of the different pulse shapes, both for single pulses and the TBS protocol, to determine the optimal current direction.

### Intermittent vs continuous TBS

Intermittent TBS is a protocol that tends to increase the excitability of the targeted area. In its counterpart, continuous (c)TBS, the 50Hz bursts are repeated continuously at 5Hz, without the 8s break between burst trains, and this has been shown to decrease excitability. While cTBS was not tested in this study, the effects of using monophasic pulses may translate to the cTBS protocol as well, as inhibitory protocols applying monophasic pulses at 1Hz have been shown to decrease the excitability more effectively than biphasic pulses [4, 8].

### Further considerations to improve TBS

Monophasic pulses approximating the pulse shape of conventional stimulators were used in this study, but other pulse shapes or widths may cause larger effects in TBS and other repetitive TMS protocols. With the newer TMS devices such as the pTMS device used in this study, researchers gain the ability to investigate more parameters to optimise the effects of TBS. Other studies looking at the number of pulses per burst [36], different stimulation intervals [34, 37] as well as the total number of pulses [24, 38] show further possible avenues to increase the plasticity effects induced by the stimulation.

This study was conducted by applying iTBS to the primary motor cortex, which is a common target to test plasticity effects of stimulation protocols. However, in clinical practice targets other than M1 are of interest. For example, repetitive TMS is used in the treatment of major depressive disorder (MDD), where it is applied daily to the dorsolateral prefrontal cortex (DLPFC) over several weeks [39]. Further studies will be required to understand how the results observed here after motor cortex stimulation may translate to other brain areas, cognitive functions and disease states.

### Vertex stimulation as control condition

In the control condition, iTBS was applied to the vertex of the head, using the same parameters as in the active conditions, to achieve a realistic control condition with similar skin sensation and audio effects [23, 40]. The purpose of the control condition was to control for the natural variability observed in MEP measurements when taken over the same time period as in the active conditions. However, as the brain was still actively stimulated in this condition, albeit in a different location, brain network effects may have influenced those results.

### Limitations

Due to the technical setup of the study, it was conducted in a single-blinded manner, where the participants were blinded to the condition and the study hypothesis, but the experimenters were aware of the stimulation condition. This was partially due to the online programming needed for the custom-made pTMS device to generate monophasic or biphasic pulses and the fact that the coil was placed in a different location during the control condition. Additionally, the test sessions were 2-3h long, during which experimenters interacted with the participants, albeit as little as possible, which may have had an influence on the results and the MEP variability, though the use of a within-participants crossover design should help to mitigate this potential issue to some extent.

## Conclusions

This study confirms that the pulse shape affects the group level plasticity effects induced after iTBS, with monophasic pulses leading to larger increases in MEP amplitude than conventional biphasic. This adds to the literature exploring improvements of the TBS protocol in the hope of enhancing plasticity induction for applications in basic neuroscience and medical practice such as depression therapy.

## Acknowledgements

This work was supported by a program grant from the MRC Brain Network Dynamics Unit at the University of Oxford (MRC MC_UU_0003/3), including an MRC iCASE studentship, and supplemental funding to TD by the Royal Academy of Engineering. JO’S is supported by a Sir Henry Dale Fellowship from the Royal Society and the Wellcome Trust (215451/Z/19/Z). MF holds a postdoctoral research fellowship from Guarantors of Brain. CJS holds a Senior Research Fellowship, funded by the Wellcome Trust (224430/Z/21/Z). This research was supported by the NIHR Oxford Health Biomedical Research Centre (NIHR203316). The views expressed are those of the authors and not necessarily those of the NIHR or the Department of Health and Social Care. The Wellcome Centre for Integrative Neuroimaging is supported by core funding from the Wellcome Trust (203139/Z/16/Z and 203139/A/16/Z). The laboratory space and equipment were supported by the University of Oxford John Fell Fund and the Wellcome Trust Institutional Strategic Support Fund (to CJS).

The authors would like to thank Magstim Company Ltd (UK) for providing the stimulation coil and valuable guidance on device design considerations. The authors would also like to thank Verena Sarrazin, Lorenzo Mazzaschi and Esraa Shaban for their help in collecting the data.

## Rights Retention

For the purpose of Open Access, the authors have applied a CC BY public copyright licence to any Author Accepted Manuscript version arising from this submission.

## Data availability statement

The data and relevant scripts will be made available on the Open Science Framework

## Authorship contributions

**Karen Wendt:** Conceptualisation, Methodology, Investigation, Formal analysis, Writing – Original Draft; **Majid Memarian Sorkhabi:** Methodology, Investigation, Resources – designed and built pTMS device, Writing - Review & Editing; **Charlotte J. Stagg:** Methodology, Writing - Review & Editing; **Melanie K Fleming:** Methodology, Writing - Review & Editing; **Timothy Denison:** Conceptualisation, Resources – designed and built pTMS device, Writing - Review & Editing, Supervision, Funding acquisition; **Jacinta O’Shea:** Conceptualisation, Methodology, Writing - Review & Editing, Supervision, Funding acquisition.

## Notes

**Conflict of interest and financial disclosures** TD, MMS and KW have pending patent applications for TMS device circuits and control algorithms. TD and KW have received grant funding (including an MRC iCASE studentship) and materials through collaboration agreements from Magstim Ltd. MMS is currently employed by Magstim Ltd. JO’S has acted as a consultant for Welcony Inc. and serves on the Scientific Advisory Board of Plato Science. CJS and MF have no conflicts of interest.

### Competing Interest Statement

TD, MMS and KW have pending patent applications for TMS device circuits and control algorithms. TD and KW have received grant funding (including an MRC iCASE studentship) and materials through collaboration agreements from Magstim Ltd. MMS is currently employed by Magstim Ltd. JOS has acted as a consultant for Welcony Inc. and serves on the Scientific Advisory Board of Plato Science. CJS and MF have no conflicts of interest.

### Summary of Updates

Increased level of detail in the methods section, Supplemental files updated

## References

[1] P. M. Rossini et al., “Non-invasive electrical and magnetic stimulation of the brain, spinal cord, roots and peripheral nerves: Basic principles and procedures for routine clinical and research application. An updated report from an I.F.C.N. Committee,” Clin Neurophysiol, vol. 126, no. 6, pp. 1071–1107, 2015.

[2] J. P. Lefaucheur et al., “Evidence-based guidelines on the therapeutic use of repetitive transcranial magnetic stimulation (rTMS): An update (2014–2018),“ vol. 131, ed: Elsevier Ireland Ltd, 2020, pp. 474–528.

[3] A. S. Aberra, B. Wang, W. M. Grill, and A. V. Peterchev, “Simulation of transcranial magnetic stimulation in head model with morphologically-realistic cortical neurons,“ Brain Stimulation, vol. 13, no. 1, pp. 175–189, 2020.

[4] M. Sommer, N. Lang, F. Tergau, and W. Paulus, “Neuronal tissue polarization induced by repetitive transcranial magnetic stimulation?,“ (in eng), Neuroreport, vol. 13, no. 6, p. 809, 2002.

[5] M. Sommer et al., “Half sine, monophasic and biphasic transcranial magnetic stimulation of the human motor cortex,“ Clin Neurophysiol, vol. 117, no. 4, pp. 838–844, 2006.

[6] N. Arai et al., “Differences in after-effect between monophasic and biphasic high-frequency rTMS of the human motor cortex,“ Clinical Neurophysiology, vol. 118, no. 10, pp. 2227–2233, 2007.

[7] R. Hannah and J. C. Rothwell, “Pulse Duration as Well as Current Direction Determines the Specificity of Transcranial Magnetic Stimulation of Motor Cortex during Contraction,“ (in eng), Brain Stimul, vol. 10, no. 1, pp. 106–115, 2017.

[8] J. L. Taylor and C. K. Loo, “Stimulus waveform influences the efficacy of repetitive transcranial magnetic stimulation,“ Journal of Affective Disorders, vol. 97, no. 1, pp. 271–276, 2007.

[9] M. MemarianSorkhabi, M. Benjaber, K. Wendt, T. West, D. Rogers, and T. Denison, “Programmable Transcranial Magnetic Stimulation-A Modulation Approach for the Generation of Controllable Magnetic Stimuli,“ IEEE Trans. on Biomedical Engineering, pp. 1–1, 2020.

[10] A. V. Peterchev and M. E. Riehl, “Transcranial Magnetic Stimulators,&; The Oxford Handbook of Transcranial Stimulation, Second Edition: Oxford University Press, p. 0. [Online]. Available: https://doi.org/10.1093/oxfordhb/9780198832256.013.3

[11] Y.-Z. Huang, M. J. Edwards, E. Rounis, K. P. Bhatia, and J. C. Rothwell, “Theta Burst Stimulation of the Human Motor Cortex,& Neuron, vol. 45, no. 2, pp. 201–206, 2005.

[12] Y.-Z. Huang et al., “Plasticity induced by non-invasive transcranial brain stimulation: A position paper,“ Clinical Neurophysiology, vol. 128, no. 11, pp. 2318–2329, 2017.

[13] K. Nakamura et al., “Variability in Response to Quadripulse Stimulation of the Motor Cortex,“ Brain Stimulation: Basic, Translational, and Clinical Research in Neuromodulation, vol. 9, no. 6, pp. 859–866, 2016.

[14] M. Hamada et al., “Bidirectional long-term motor cortical plasticity and metaplasticity induced by quadripulse transcranial magnetic stimulation,“ J Physiol, vol. 586, no. 16, pp. 3927–3947, 2008.

[15] A. V. Peterchev, K. Dʼ Ostilio, J. C. Rothwell, and D. L. Murphy, “Controllable pulse parameter transcranial magnetic stimulator with enhanced circuit topology and pulse shaping,“ Journal of Neural Engineering, vol. 11, no. 5, p. 056023, 2014.

[16] Z. Zeng, L. M. Koponen, R. Hamdan, Z. Li, S. M. Goetz, and A. V. Peterchev, “Modular multilevel TMS device with wide output range and ultrabrief pulse capability for sound reduction,“ Journal of Neural Engineering, vol. 19, no. 2, p. 026008, 2022.

[17] J. O. Nieminen et al., “Multi-locus transcranial magnetic stimulation system for electronically targeted brain stimulation,“ Brain Stimulation, vol. 15, no. 1, pp. 116–124, 2022.

[18] K. Wendt, M. M. Sorkhabi, J. O’Shea, H. Cagnan, and T. Denison, “Comparison between the modelled response of primary motor cortex neurons to pulse-width modulated and conventional TMS stimuli,“ (in eng), Annu Int Conf IEEE Eng Med Biol Soc, vol. 2021, pp. 6058–6061, 2021.

[19] M. Memarian Sorkhabi, K. Wendt, J. O’Shea, and T. Denison, “Pulse width modulation-based TMS: Primary motor cortex responses compared to conventional monophasic stimuli,“ Brain Stimulation: Basic, Translational, and Clinical Research in Neuromodulation, vol. 15, no. 4, pp. 980–983, 2022.

[20] R. C. Oldfield, “The assessment and analysis of handedness: the Edinburgh inventory,“ (in eng), Neuropsychologia, vol. 9, no. 1, pp. 97–113, 1971.

[21] S. Rossi, M. Hallett, P. M. Rossini, and A. Pascual-Leone, “Safety, ethical considerations, and application guidelines for the use of transcranial magnetic stimulation in clinical practice and research,“ Clinical Neurophysiology, vol. 120, no. 12, pp. 2008–2039, 2009.

[22] R. Gentner, K. Wankerl, C. Reinsberger, D. Zeller, and J. Classen, “Depression of Human Corticospinal Excitability Induced by Magnetic Theta-burst Stimulation: Evidence of Rapid Polarity-Reversing Metaplasticity,“ Cerebral Cortex, vol. 18, no. 9, pp. 2046–2053, 2008.

[23] C. Nettekoven et al., “Dose-dependent effects of theta burst rTMS on cortical excitability and resting-state connectivity of the human motor system,” (in eng), J Neurosci, vol. 34, no. 20, pp. 6849–59, 2014.

[24] D. M. McCalley, D. H. Lench, J. D. Doolittle, J. P. Imperatore, M. Hoffman, and C. A. Hanlon, “Determining the optimal pulse number for theta burst induced change in cortical excitability,“ Scientific Reports, vol. 11, no. 1, p. 8726, 2021.

[25] D. Lakens, “Calculating and reporting effect sizes to facilitate cumulative science: a practical primer for t-tests and ANOVAs,“ Frontiers in Psychology, Review vol. 4, 2013.

[26] D. Curran-Everett and C. L. Williams, “Explorations in statistics: the analysis of change,“ Advances in Physiology Education, vol. 39, no. 2, pp. 49–54, 2015.

[27] A. Kuznetsova, P. B. Brockhoff, and R. H. B. Christensen, “lmerTest Package: Tests in Linear Mixed Effects Models,“ Journal of Statistical Software, vol. 82, no. 13, pp. 1 – 26, 2017.

[28] D. Bates, M. Mächler, B. Bolker, and S. Walker, “Fitting Linear Mixed-Effects Models Using lme4,“ Journal of Statistical Software, vol. 67, no. 1, pp. 1 – 48, 2015.

[29] A. Jannati, G. Block, L. M. Oberman, A. Rotenberg, and A. Pascual-Leone, “Interindividual variability in response to continuous theta-burst stimulation in healthy adults,“ Clinical Neurophysiology, vol. 128, no. 11, pp. 2268–2278, 2017.

[30] L. Schilberg, T. Schuhmann, and A. T. Sack, “Interindividual Variability and Intraindividual Reliability of Intermittent Theta Burst Stimulation-induced Neuroplasticity Mechanisms in the Healthy Brain,“ Journal of Cognitive Neuroscience, vol. 29, no. 6, pp. 1022–1032, 2017.

[31] Y.-Z. Huang, J. C. Rothwell, M. J. Edwards, and R.-S. Chen, “Effect of Physiological Activity on an NMDA-Dependent Form of Cortical Plasticity in Human,“ Cerebral Cortex, vol. 18, no. 3, pp. 563–570, 2008.

[32] D. Burke and E. Pierrot-Deseilligny, “Caveats when studying motor cortex excitability and the cortical control of movement using transcranial magnetic stimulation,“ Clinical Neurophysiology, vol. 121, no. 2, pp. 121–123, 2010.

[33] A. M. Vallence, M. R. Goldsworthy, N. A. Hodyl, J. G. Semmler, J. B. Pitcher, and M. C. Ridding, “Inter- and intra-subject variability of motor cortex plasticity following continuous theta-burst stimulation,“ Neuroscience, vol. 304, pp. 266–278, 2015.

[34] N. Y. Tse et al., “The effect of stimulation interval on plasticity following repeated blocks of intermittent theta burst stimulation,“ Scientific Reports, vol. 8, no. 1, p. 8526, 2018.

[35] C. Mastroeni et al., “Brain-derived neurotrophic factor--a major player in stimulation-induced homeostatic metaplasticity of human motor cortex?,” (in eng), PLoS One, vol. 8, no. 2, p. e57957, 2013.

[36] Q. Meng et al., “A high-density theta burst paradigm enhances the aftereffects of transcranial magnetic stimulation: Evidence from focal stimulation of rat motor cortex,“ Brain Stimulation: Basic, Translational, and Clinical Research in Neuromodulation, vol. 15, no. 3, pp. 833–842, 2022.

[37] C. Abboud Chalhoub, S. Awasthi, S. Exley, B. Luber, S. Lisanby, and Z.-D. Deng, “Optimizing the Effects of Theta-Burst Stimulation (TBS),“ Biological Psychiatry, vol. 87, no. 9, Supplement, p. S317, 2020.

[38] O. L. Gamboa, A. Antal, V. Moliadze, and W. Paulus, “Simply longer is not better: reversal of theta burst after-effect with prolonged stimulation,“ Experimental Brain Research, vol. 204, no. 2, pp. 181–187, 2010.

[39] D. M. Blumberger et al., “Effectiveness of theta burst versus high-frequency repetitive transcranial magnetic stimulation in patients with depression (THREE-D): a randomised noninferiority trial,“ The Lancet, vol. 391, no. 10131, pp. 1683–1692, 2018.

[40] U. Herwig, L. Cardenas-Morales, B. J. Connemann, T. Kammer, and C. Schönfeldt-Lecuona, “Sham or real—Post hoc estimation of stimulation condition in a randomized transcranial magnetic stimulation trial,“ Neuroscience Letters, vol. 471, no. 1, pp. 30–33, 2010.

