## Supplementary Figures for "The effect of pulse shape in theta-burst stimulation: monophasic vs biphasic TMS"

### Supplementary File

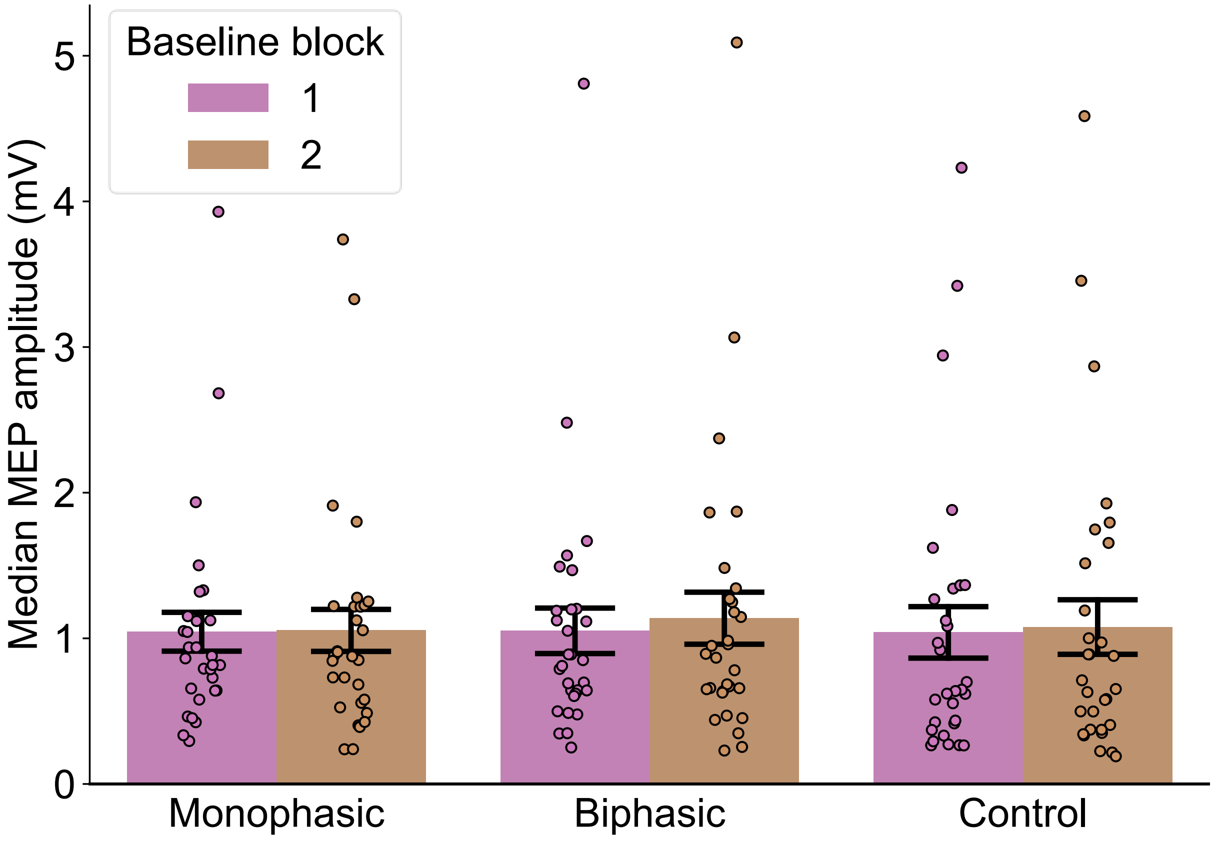


*Fig. S1: Group mean of the median peak-to-peak MEP amplitude for each of the two baseline blocks averaged across participants in each condition. Each baseline block consisted of MEPs elicited by 30 single TMS pulses at 120% of the resting motor threshold. The group mean MEP amplitude was close to 1 mV for each condition and block, but the median amplitudes of individual participants varied. Individual participants medians are indicated by dots, the bars indicate group means and the error bars represent +/- 1 standard error of the mean.*

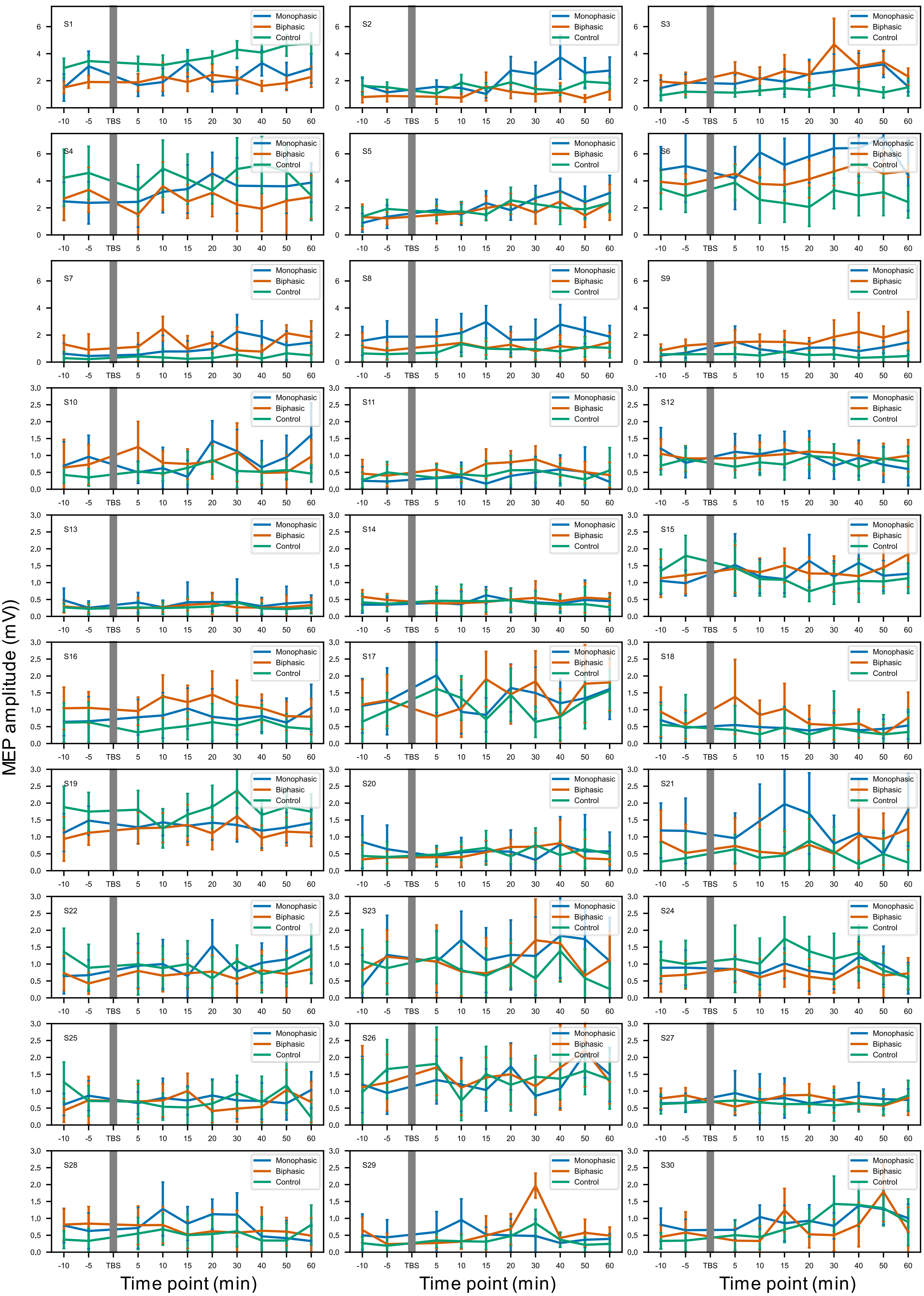
S1
S2
S3
S4
S5
S6
S7
S8
S9
S10
S11
S12
S13
S14
S15
S16
S17
S18
S19
S20
S21
S22
S23
S24
S25
S26
S27
S28
S29
S30
Monophasic
Biphasic
Control
MEP amplitude (mV)
Time point (min)
-10
-5
TBS
5
10
15
20
30
40
50
60

*Fig. S2: Median block-wise MEP amplitudes for all participants and TMS conditions. Each plot shows the data of one participant across all data collection time points in all TBS conditions in mV. The error bars represent the standard deviation. Note: the y-axis range has the same scale for S1-S9, and a different range for S10-S30.*

**Electromyography setup**

Surface electromyography (EMG) of the right FDI was recorded using disposable neonatal ECG electrodes (Kendall, Cardinal Health, UK) in a belly-tendon montage with a ground electrode over the ulnar styloid process. EMG signals were sampled at 10 kHz, amplified with a gain of 1000, filtered (10 Hz - 1000 Hz) and recorded using a D440 Isolated Amplifier (Digitimer, Welwyn Garden City, UK), a Micro1401 (Cambridge Electronic Design, Cambridge, UK) and Signal software version 7.01 (Cambridge Electronic Design). The unwanted 50 Hz line noise was attenuated by subtracting a replica of the measured noise from the input signal using a HumBug Noise Eliminator (Digitimer).
